# Two-dimensional local Fourier image reconstruction via domain decomposition Fourier continuation method

**DOI:** 10.1101/322438

**Authors:** Ruonan Shi, Jae-Hun Jung, Ferdinand Schweser

**Affiliations:** Department of Mathematics, State University of New York at Buffalo, Buffalo, NY 14260, U.S.A.; Department of Data Science, Ajou University, Yeongtung-gu, Suwon 16499, Korea; Buffalo Neuroimaging Analysis Center, Department of Neurology, Jacobs School of Medicine and Biomedical Sciences at the University at Buffalo, Buffalo, NY 14260, U.S.A.; Center for Biomedical Imaging, Clinical and Translational Science Institute, University at Buffalo, Buffalo, NY 14260, USA

**Author notes:** (Ruonan Shi), (Jae-Hun Jung), (Ferdinand Schweser).

## Abstract

The MRI image is obtained in the spatial domain from the given Fourier coefficients in the frequency domain. It is costly to obtain the high resolution image because it requires higher frequency Fourier data while the lower frequency Fourier data is less costly and effective if the image is smooth. However, the Gibbs ringing, if existent, prevails with the lower frequency Fourier data. We propose an efficient and accurate local reconstruction method with the lower frequency Fourier data that yields sharp image profile near the local edge. The proposed method utilizes only the small number of image data in the local area. Thus the method is efficient. Furthermore the method is accurate because it minimizes the global effects on the reconstruction near the weak edges shown in many other global methods for which all the image data is used for the reconstruction. To utilize the Fourier method locally based on the local non-periodic data, the proposed method is based on the Fourier continuation method. This work is an extension of our previous 1D Fourier domain decomposition method to 2D Fourier data. The proposed method first divides the MRI image in the spatial domain into many subdomains and applies the Fourier continuation method for the smooth periodic extension of the subdomain of interest. Then the proposed method reconstructs the local image based on *L*_2_ minimization regularized by the *L*_1_ norm of edge sparsity to sharpen the image near edges. Our numerical results suggest that the proposed method should be utilized in dimension-by-dimension manner instead of in a global manner for both the quality of the reconstruction and computational efficiency. The numerical results show that the proposed method is effective when the local reconstruction is sought and that the solution is free of Gibbs oscillations.

## Introduction

Magnetic resonance imaging (MRI) is a commonly used medical imaging technique. MRI image reconstruction is based on the inverse Fourier transform of a frequency-limited acquired Fourier spectrum of the object. In this work, we are interested in the 2D Fourier reconstruction of given MRI data. Before we continue to the 2D MRI image reconstruction, we should start from the basic 1D Fourier reconstruction. For the 1D Fourier reconstruction, we consider the problem of how to reconstruct a piecewise smooth function *f*(*x*) : Ω → ℝ, Ω = [−1,1] on a uniform grid *x_j_* = −1 + *j*Δ*x*, *j* = 0,1, ···, 2*N*, 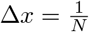 when its Fourier coefficients (also known as k-space data in MRI), 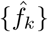, are given. The Fourier coefficients of *f*(*x*) are defined by

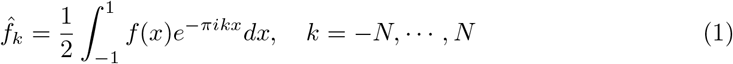

and the Fourier partial sum *f*_*N*_(*x*), which is the image space reconstruction commonly obtained in MRI, is

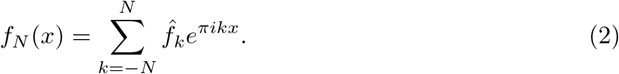

Since the function that we want to find, *f*(*x*), is a piecewise smooth function, the approximation by Eq (2) may yield the Gibbs phenomenon.

When the patient’s MRI image is obtained, usually we want to reconstruct the image so as to reveal the detailed structure of a particular region with the given Fourier coefficients. However, most reconstruction methods are carried out in a global manner, so they are computationally expensive. And since global methods provide the reconstruction in one piece, so the quality of the reconstruction near different edges is not equal([12] shows this limitation). In this work, we propose a local method that focuses on and yields the local reconstruction of the subdomain such that an accurate and non-oscillatory sharp image reconstruction is achieved in that subdomain. If we patch all these local reconstructions together to constitute the whole image, the result will be more accurate than the one obtained by the global method while the subdomain that we are interested in is much enhanced.

As an example, we propose a local method base on the *L*_1_ regularization method proposed in [13]. The method by [13], known as the sparse polynomial annihilation (PA) method, is an *L*_1_ regularization method based on sparsity of edges for the Fourier reconstruction, for obtaining the sharp image profiles near edges. By using the edge sparsity in the *L*_1_ regularization term, this approach can reduce the Gibbs oscillations near edges and provide a sharp reconstruction. However, as other reconstruction methods, this approach is a global method, which intrinsically puts more weight on strong edges (jumps with large magnitudes) than on weak edges (jumps with small magnitudes). This can sharpen the reconstruction near strong edges, but the reconstruction near weak edges often becomes similar to or even worse than the original Fourier reconstruction. Thus, if the area we are interested in is near weak edges, this approach may not be effective. If the *L*_1_ regularization can be done locally, the reconstruction near weak edges can be improved. Such local operation could be done by our previous 1D method of domain decomposition Fourier *L*_2_ and *L*_1_ minimization based on the Fourier continuation [12]. The key element of our local method is to split the given domain into multiple subdomains first and then carry out the *L*_1_ minimization individually in each subdomain. If necessary, we patch all the reconstruction in each subdomain together.

In this paper, we extend our previous 1D domain decomposition Fourier reconstruction method to 2D image reconstruction. We propose the following 2D methods, 1) the global 2D Fourier continuation sparse PA method and 2) the dimension by dimension Fourier continuation sparse PA method. The dimension by dimension is basically a 1D Fourier continuation applied in *x*− and *y*−directions separately. By comparing numerically the global 2D Fourier continuation sparse PA method with the dimension by dimension Fourier continuation sparse PA method, we show that the dimension by dimension method should be used for both accuracy and computational efficiency. In this paper, we also provide empirically the best range of the control value of the *L*_1_ regularization, λ, to use. Our method is efficient and flexible in the sense that we can stitch all the local reconstructions from each subdomain to form the accurate whole complete image and that the constituted one is more accurate than the image by the global method. In this paper we provide detailed numerical results using the Shepp-Logan phantom image and the real MRI data.

This paper is composed of the following sections. In Section 2, a brief explanation of the 1D domain decomposition Fourier continuation method based on the edge sparsity proposed in [12] is given. In Section 3, the extension of the 1D method to the 2D Fourier reconstruction will be introduced. Various numerical examples are given in Section 4. Finally in Section 5, a concluding remark will be provided.

## 1D domain decomposition Fourier continuation sparse PA method

### 1D Fourier continuation

We briefly explain two approaches of the Fourier continuation method in 1D. More detailed explanations can be found in [4–8,10,11].

### Global Fourier continuation method

Suppose that we have a non-periodic function *f* : *I*_0_ → ℝ where *I*_0_ = [*a, b*]. Let *L* = *b*−*a*. The point values of *f, f* (*x_j_*), are given at *x_j_* = *a* + *j*Δ*x*, where 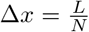 *j* = 0, ···, *N*. With the global 1D Fourier continuation method, the non-periodic function *f*(*x*) is extended to a periodic function *g*(*z*) over *I*_2_ = [*a, b + d*] (where *d* = γΔ*x*, γ is a positive integer) with periodicity of *L + d*. The positive value of γ is arbitrary and the optimal value of γ is function dependent. Practically the value of γ is chosen empirically in consideration of computational efficiency.

The periodic extension *g*(*z*) can be written as the following Fourier sum since *g*(*z*) is periodic over *I*_2_

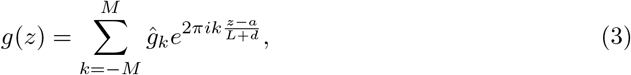

where *M* is a nonnegative integer. The unknown coefficients *ĝ_k_* can be obtained by matching *g*(*z*) with *f*(*x*) at *z* = *x = x_j_* such that

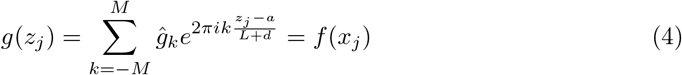

where *z_j_* = *a* + *j*Δ*x* = *x_j_*, *j* = 0,1, ···, *N*. Let the element of the coefficient matrix *A* be defined by

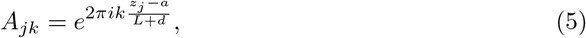

and let 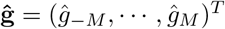 and **f** = (*f* (*x_0_*), ···, *f*(*x_N_*))*^T^*. Then *ĝ_k_* in Eq (3) can be found by solving the following linear system

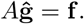

If *M = N*/2, *ĝ_k_* are uniquely determined. If *M ≠ N*/2, the system can be solved by using the pseudo inverse based on the singular value decomposition (SVD),

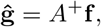

where *A*_+_ denotes the pseudo inverse of *A*. The matrix *A* easily becomes ill-posed with the large values of *N* and *M*, which leads us to the method in the following section.

### Fourier continuation method using the boundary values

The global Fourier continuation method is simple, but since this method uses every *f*(*x_i_*) to find the extended periodic function, the reconstruction is easily affected by round-off errors and the Runge phenomenon still exists if *f*(*x*) is a Runge function. To minimize such issues, the local Fourier continuation method [10,11] has been used which utilizes *f*(*x_i_*) near the domain boundaries only. We briefly explain this local Fourier continuation method below.

The extended periodic function *g*(*z*) over [*a, b* + *d*] is defined as

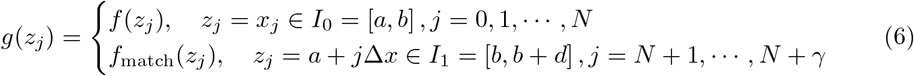

where *f*_match_(*z*) is called the matching function. Here, *g*(*z*) is defined different from *g*(*z*) in Eq (3). The local Fourier continuation method uses only local values near the boundaries to find the matching function. Define the intervals *I*_left_ = [*b −δ, b*] and *I*_right_ = [*b + d, b + d + δ*] (where *δ* = *β*Δ*x, d* = *γ*Δ*x*, *β* and γ are positive integers). With the values of *f*(*x*) over *I*_left_ and *I*_right_, the matching function f_match_(z) over these intervals is defined by

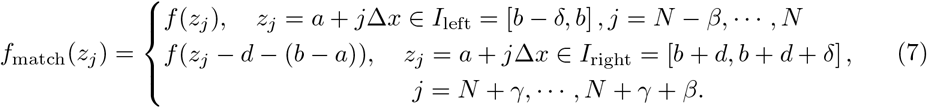

*f*_match_ (*z*) is a periodic function defined over *I*_3_ = [*b* − δ, *b* + 2*d* + δ] with the periodicity of 2(*d* + δ). By finding *f*_match_, the extended periodic function *g*(*z*) in Eq (6) is found. To find *f*_match_(*z*) over the whole interval *I*_3_, we use following formula

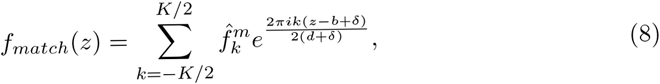

where 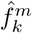 are the unknown Fourier coefficients of *f*_match_(*z*). We find 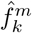 by matching the matching function with the given function. That is,

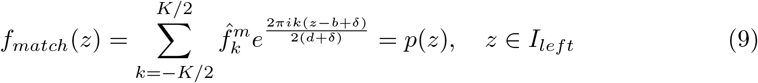

and

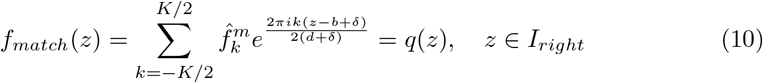

at *Q* distinct points in each interval. *K* is an integer that *K* < *Q*. Here *p*(*z*) and *q*(*z*) are found by using (*β* + 1) grid points such that the degree of *p*(*z*) and *q*(*z*) is less than or equal to *β*. If the degree is *β*, then *p*(*z*) and *q*(*z*) are the local interpolation of *f*(*x*) over *I*_left_ and *I*_right_, respectively. If the degree is less than *β*, then *p*(*z*) and *q*(*z*) can be found by solving the over-determined system in the least-squares sense. To minimize round-off errors, people usually use the Gram polynomial for **p*(*z*)* and *q*(*z*) and construct the linear system [10]. By solving Eq (9) and Eq (10) in the least-squares sense using SVD we find 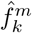. Then using Eq (8) and Eq (6), we find the matching function *f*_match_(*z_j_*),*j* = *N*, ···, *N* + γ and obtain the desired extended function *g*(*z_j_*),*j* = 1 ···, *N* + γ.

### Convex optimization: *L*_1_ regularization based on sparsity of edges

The following convex optimization problem using *L*_1_ regularization based on sparsity of edges was proposed in [13], which yields sharp Fourier reconstruction near edges,

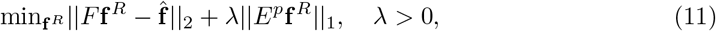

where || · ||_2_ and || · ||_1_ denote the vector *L*_2_ and *L*_1_ norms, respectively. The first term is the fidelity term. **f**^R^ is the reconstruction we want to find and **f̂** is the given Fourier coefficient vector. The edge operator *E^p^* is the sparse polynomial annihilation transform and the superscript *p* denotes the order of the derivative of the interpolation [2, 3]. The polynomial annihilation (PA) is basically higher order derivative of the interpolation. Thus *E^p^***f***^R^* has large values at the discontinuities but vanishes in the smooth regions of the function. In such a way, *E^p^***f**^*R*^ represents sparsity. The constant λ > 0 is a free parameter whose optimal value is chosen empirically [13]. In [13] it was shown that the *L*_1_ regularization with the sparse PA method yields a better reconstruction than the filtering or TV regularization.

### 1D domain decomposition Fourier continuation sparse PA method

In [12] the domain decomposition sparse PA method was proposed, with which the given domain is split into multiple subdomains and the sparse PA method is applied separately in each individual domain. To make the minimization of Eq (11) done locally, we used the Fourier continuation method.

**Remark 1** *Here we should note that our domain decomposition Fourier continuation method is not limited to the sparse PA method. The *L*_1_ regularization term in Eq (11) can be replaced with other terms such as TV norm*.

For the domain decomposition Fourier continuation method, we split the domain into multiple subdomains {[*x*_0_,*x*_*j*_1__], [*x*_*j*_1__,*x*_*j*_2__], ···, [*x*_*j*_*k*-1__, *x*_*j*_k__], [*x*_*j*_k__,*x*_2*N*_]}, 0 < *j*_1_ < *j*_2_ < ··· < *j_k_* < 2*N* and carry out the minimization for each subdomain separately. For example, suppose that we reconstruct the function on the subdomain [*x*_*j*1_, *x*_*j*2_]. The function is not necessarily periodic in this subdomain. To deal with this non-periodic function with the Fourier method, we use the Fourier continuation method. However, as shown in [12], the direct application of the Fourier continuation method to the subdomain [*x*_*j*_1__, *x*_*j*_2__] causes some oscillations near the domain boundaries when trying to stitch the all the reconstructions from each subdomain. To avoid such oscillations and apply the Fourier continuation successfully, we first extend the subdomain so that the extended domain contains [*x*_*j*_1__, *x*_*j*_2__]. The easiest way to find this extension is to add a fixed number of extra points to both boundaries. A more sophisticated way is presented in our previous work [12] based on the TV analysis. Let *n_m_* be a fixed integer that determines how many extra points are added at each boundary. Thus the extension becomes 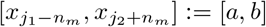, which contains *K*_0_ = (*j*_2_ + *n_m_*) − (*j*_1_ − *n_m_*) + 1 uniform grid points. Thus we have the non-periodic data *f_N_*(*x*)(obtained from all Fourier coefficients via Eq. 2) on *K_0_* uniform grid points over [*a, b*].

Then we use the following convex optimization method in this subdomain using *L*_1_ regularization based on the sparsity of edges

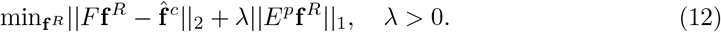

*F* is a transform matrix explained below. 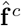 is a Fourier coefficient vector which needs to be found before solving Eq (12). This 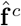 is not the one defined by Eq (1), but is obtained from *f_N_*(*x*) over [*a, b*]. For the extended subdomain, [*a, b*] we apply the Fourier continuation method to *f_N_*(*x*) over [*a, b*] [12] and obtain the extended periodic function *f_ex_*(*x*) over the extended interval [*a, b + d*], where *d* = *γ*Δ*x*. γ is a fixed integer that determines the number of points on the extended interval as defined in the previous section. Let *K* = *K*_0_ + γ. With the *f_ex_*(*x*) over the extended interval [*a, b + d*], the corresponding Fourier coefficients, 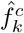, can be found by using the discrete Fourier transform [9],

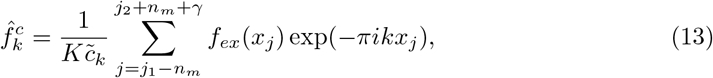

where

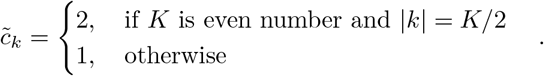

Let 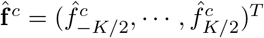 and *F* be the transform matrix whose elements are *F_kj_* = exp(−*ikπx_j_*)/*Kc̃_k_* from Eq (13).

The optimization problem Eq (12) is solved using the Matlab CVX package [1]. f^*R*^ is the reconstruction we get over extended interval [*a, b + d*]. The reconstruction by Eq (12) is over the domain [*a, b + d*]. We only take the reconstruction over the interval [*x*_*j*_1__, *x*_*j*_2__] ⊂ [*a, b*] ⊂ [*a, b + d*].

### 2D domain decomposition Fourier continuation method

Suppose that we have a periodic function *f*_0_ : *J*_0_ → ℝ over *J*_0_ = [*a*_0_, *b*_0_]^2^. Let *L*_0_ = *b*_0_−*a*_0_. Further suppose that the values of *f*_0_ are given at a set of uniform grid points, (*x_i_*, *y_j_*), *x_i_* = *a*_0_ + *i*Δ and *y_j_* = *a*_0_ + *j*Δ, where 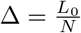 and *i* = 0, ···, *N, j* = 0, ···, *N*. We consider the problem of how to reconstruct the 2D image on *n* × *n* uniform grid points over *J* = [*a, b*] × [*c, e*] ⊂ *J*_0_, where

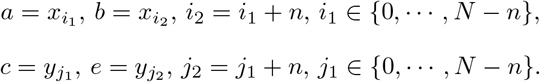

In practice, the subdomain size is determined by the size of the region where the local Gibbs oscillations are dominant. Since the function *f*_0_ over the subdomain *J* is non-periodic in general and the Fourier coefficients computed based on the function values within *J* are Gibbs-contaminated, we apply the Fourier continuation method to *J*. Before we use the Fourier continuation method to reduce the Gibbs oscillations near the boundaries of *J* we first need the function values on a larger subdomain than *J* as in the 1D case. Here we choose the same number of extra grid points to add to all four boundaries, *n_m_*. Then we have the new subdomain *J*_1_ = [*a*_1_, *b*_1_] × [*c*_1_, *e*_1_], where

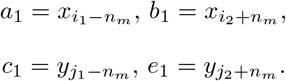

Thus over this new subdomain *J*_1_ we have a new non-periodic function *f* : *J*_1_ → ℝ, that *f*(*x_i_*, *y_j_*) = *f*_0_(*x_i_*, *y_j_*), where *i* = *i*_1_−*n_m_*, *i*_1_−*n_m_* + 1, ···, *i*_2_ + *n_m_* and *j* = *j*_1_−*n_m_*, *j_i_*−*n_m_* + 1, ···, *j*_2_ + *n_m_*. Then we can apply the following two 2D Fourier continuation sparse PA methods on function *f* on (*n* + 2*n_m_*) × (*n* + 2*n_m_*) uniform grid points over *J*_1_.

### Global 2D Fourier continuation sparse PA method

Now we have a non-periodic function *f* : *J*_1_ → ℝ over *J*_1_ = [*a*_1_, *b*_1_] × [*c*_1_, *e*_1_]. Let *L* = *b*_1_−*a*_1_ = *e*_1_−*c*_1_. And the values of *f* are given at a set of uniform grid points, (*x_i_*, *y_j_*), where *i* = *i*_1_−*n_m_*, *i*_1_−*n_m_* + 1, ···, *i*_2_ + *n_m_* and *j* = *j*_1_−*n_m_*, *j*_1_−*n_m_* + 1, ···, *j*_2_ + *n_m_*. With the global Fourier continuation method, *f*(*x,y*) is extended to a periodic function *g*(*z*^(1)^,*z*^(2)^) defined over *J*_2_ = [*a*_1_, *b*_1_ + *d*] × [*c*_1_, *e*_1_ + *d*] (where *d* = γΔ, γ is a positive integer), with periodicity of *L* + *d* on *x* and *y* directions. The periodic extension *g*(*z*^(1)^, *z*^(2)^) can be obtained as below

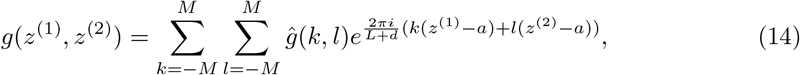

where *M* = (*n* + 2*n_m_* + γ)/2. The unknown coefficients *ĝ*(*k, l*) are determined by matching *g*(*z*^(1)^,*z*^(2)^) with *f*(*x,y*) at *z*^(1)^ = *x* = *x_i_*,*z*^(2)^ = *y* = *y_j_* such that

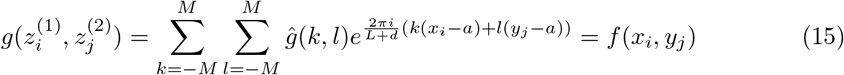

where *i* = *i*_1_−*n_m_*, *i*_1_−*n_m_* + 1, ···, *i*_2_ + *n_m_* and *j* = *j_i_*−*n_m_*, *j*_1_−*n_m_* + 1, ···, *j*_2_ + *n_m_*. The reconstruction of *f*(*x,y*) within *J*_1_ based on {*f*(*x_i_*,*y_j_*)} is given by *g*(*z*^(1)^, *z*^(2)^) for *z*^(1)^ ∈ [*a*_1_,*b*_1_] and *z*^(2)^ ∈ [*c*_1_,*e*_1_].

Then we seek **f***^R^* = {*f^R^*(*x_i_*,*y_j_*) : *i* = *i*_1_−*n_m_*, *i*_1_−*n_m_* + 1/2, *i*_1_−*n_m_* + 1, ···, *i*_2_ + *n_m_* + γ, *j* = *j*_1_−*n_m_*,*j*_1_−*n_m_* + 1/2, *j*_1_−*n_m_* + 1, ···, j_2_ + *n_m_* + γ}(a higher number of grid points is used here) by solving the convex optimization problem

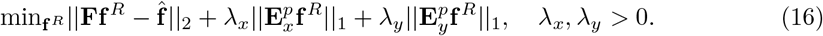

Here **f̂** is zero padding of *ĝ*(*k, l*), that

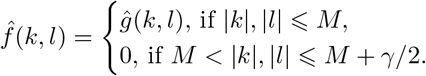

**F** is the analogous 2D Fourier transform matrix of (**??**). 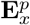 corresponds to *E_p_* in (**??**) in the *x* direction for each fixed *y_j_*, *j* = 0, ···, *N*, and 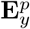 is similarly calculated for the γ direction. Finally we obtain the 2D Fourier reconstruction image, *f*(*x_i_*,*y_j_*) = *f^R^*(*x_i_*,*y^j^*), *i* = *i*_1_, *i*_1_ + 1/2, *i*_1_ + 1 ···, *i*_2_, *j* = *j*_1_, *j*_1_ + 1/2, *j*_1_ + 1, ···, *j*_2_ over the smaller domain *J*.

### Dimension by dimension Fourier continuation sparse PA method

The global approach in the previous subsection is slow. Thus we propose to use the dimension by dimension approach. That is, we apply the 1D Fourier continuation method using subdomain boundary values for each fixed *x_i_*, *i* = *i*_1_−*n_m_*, ···, *i*_2_ + *n_m_* in *x*-direction and vice versa. The non-periodic function *f*(*x_i_,y*) on {(*x_i_, y_j_*) | *j* = *j*_1_−*n_m_*, *j*_2_−*n_m_* + 1, ···, *j*_2_ + *n_m_*} is extended to a periodic function *g^R^*(*x_i_, z*) on a high resolution grid set {(*x_i_, z_I_*) | *I* = *j*_1_−*n_m_*, *j*_1_−*n_m_* + 1/2,*j*_1_−*n_m_* + 1, ···, *j*_2_ + *n_m_* + γ−1/2,*j*_2_ + *n_m_*, + γ,γ is a positive integer}, where if *I* = *j*_1_−*n_m_* + 1/2 then we have 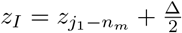. Then the corresponding Fourier coefficients, 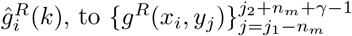 can be found using the discrete Fourier transform

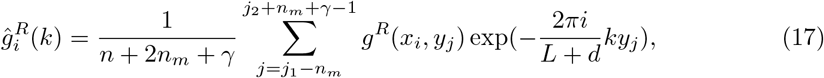

where *k* = −*M*, ···, *M*, *M* = (*n* + 2*n_m_* + γ)/2. Then we use the convex optimization method using *L*_1_ regularization based on the sparsity of edges

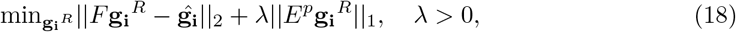

to find the reconstruction of the function, 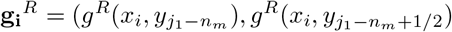, 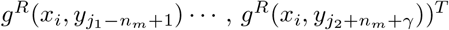, where *i* = *i*_1_ − *n_m_*, ···, *i*_2_ + *n_m_*, on 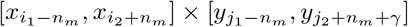. *E^p^* is the sparse PA transform matrix as (??). F is the transform matrix whose elements are

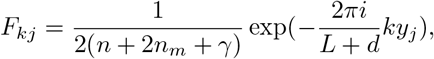

where
*k* = −*M*, ···, *M*, *j* = *j*_1_−*n_m_*, *j*_1_−*n_m_* + 1/2, *j*_1_−*n_m_*, + 1, ···, *j*_2_ + *n_m_*, + γ −1/2,*j*_2_ + *n_m_*, + γ. We update the values of f on (*n* + 2*n_m_*) × (2*n* + 4*n_m_*) grid points over *J*_1_ with these reconstruction data by letting

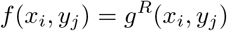

where
*i* = *i*_1_−*n_m_*, *i*_1_ − *n_m_* + 1/2,*i*_1_−*n_m_* + 1 ···, *i*_2_ + *n_m_*, *j* = *j*_1_−*n_m_*, *j*_1_−*n_m_* + 1, ···, *j*_2_ + *n_m_*.

We repeat the same procedure for *y*-direction and update the values of *f* over *J*_1_ again. We have *f*(*x, y*) on (2*n* + 4*n_m_*) × (2*n* + 4*n_m_*) uniform grid points, (*x_i_*,*y_j_*), where *i* = *i*_1_ − *n_m_*,*i*_1_ − *n_m_* + 1/2, *i*_1_ − *n_m_* + 1, ···, *i*_2_ + *n_m_*, *j* − *j*_1_ − *n_m_*, *j*_1_ − *n_m_* + 1/2, *j*_1_ − *n_m_* + 1, ···, *j*_2_ + *n_m_*. Finally we obtain the 2D Fourier reconstruction, *f*(*x_i_*,*y_j_*), *i* = *i*_1_, *i*_1_ + 1/2, *i*_1_ + 1, ···, *i*_2_, *j* = *j*_1_, *j*_1_ + 1/2, *j*_1_ + 1, ···, *j*_2_ over the smaller domain *J*.

## Numerical results

In this section, we provide various numerical examples.

### Example 1: Shepp-Logan phantom

With this example, we apply both the global 2D Fourier continuation sparse PA method and the dimension by dimension Fourier continuation sparse PA method for the reconstruction of the Shepp-Logan phantom image. This reconstructed Shepp-Logan phantom image is found by the following steps. First we use the Shepp-Logan phantom image on the 801 × 801 grid to find its 2D Fourier coefficients, 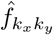, *k_x_*, *k_y_* = −400, ···, 400. We use these Fourier coefficients as the *exact* Fourier coefficients of the Shepp-Logan phantom. Using only a partial number of Fourier coefficients, i.e. *k_x_*, *k_y_* = −*N*, ···, *N*, *N* = 42 in this work, we reconstruct the Shepp-Logan phantom via DFT, *f*_*N*_, with which the 2D Fourier continuation method is applied. This experiment mimics an MRI-like acquisition with limited spatial resolution. We compare the results by both the global and dimension by dimension methods.

#### Global method

In Fig 1, the left figure on the top row is the illustration of the Shepp-Logan image on 85 × 85 grids. In the top right figure we choose two sample regions of the reconstructed Shepp-Logan, *f_N_* sample region 1 and sample region 2 in the left and right boxes respectively. We apply the global 2D Fourier continuation sparse PA method on these regions. Here we choose λ = 0.01 for both region. The best value of λ, λ = 0.01 was chosen based on the experiments. The regions in the rectangles with the solid border, *J*, are the ones where we want to find the reconstruction and the regions in the rectangles with the dashed border, *J*_1_, are the extended regions for the Fourier continuation. The subfigures in middle column of second and third row) are obtained by following procedure. First we have the 85 × 85 Fourier coefficients that were used to generate the subfigure top right, then apply zero-padding to these Fourier coefficients to generate a 169 × 169 new Fourier coefficients. By applying inverse Fourier transform of these new Fourier coefficients, we create a 169 × 169 image. Finally we take the corresponding sample region to the sample region 1 and 2 in top right. These “double resolution” images have the same matrix size as the reconstructions on the right.

**Fig 1.**
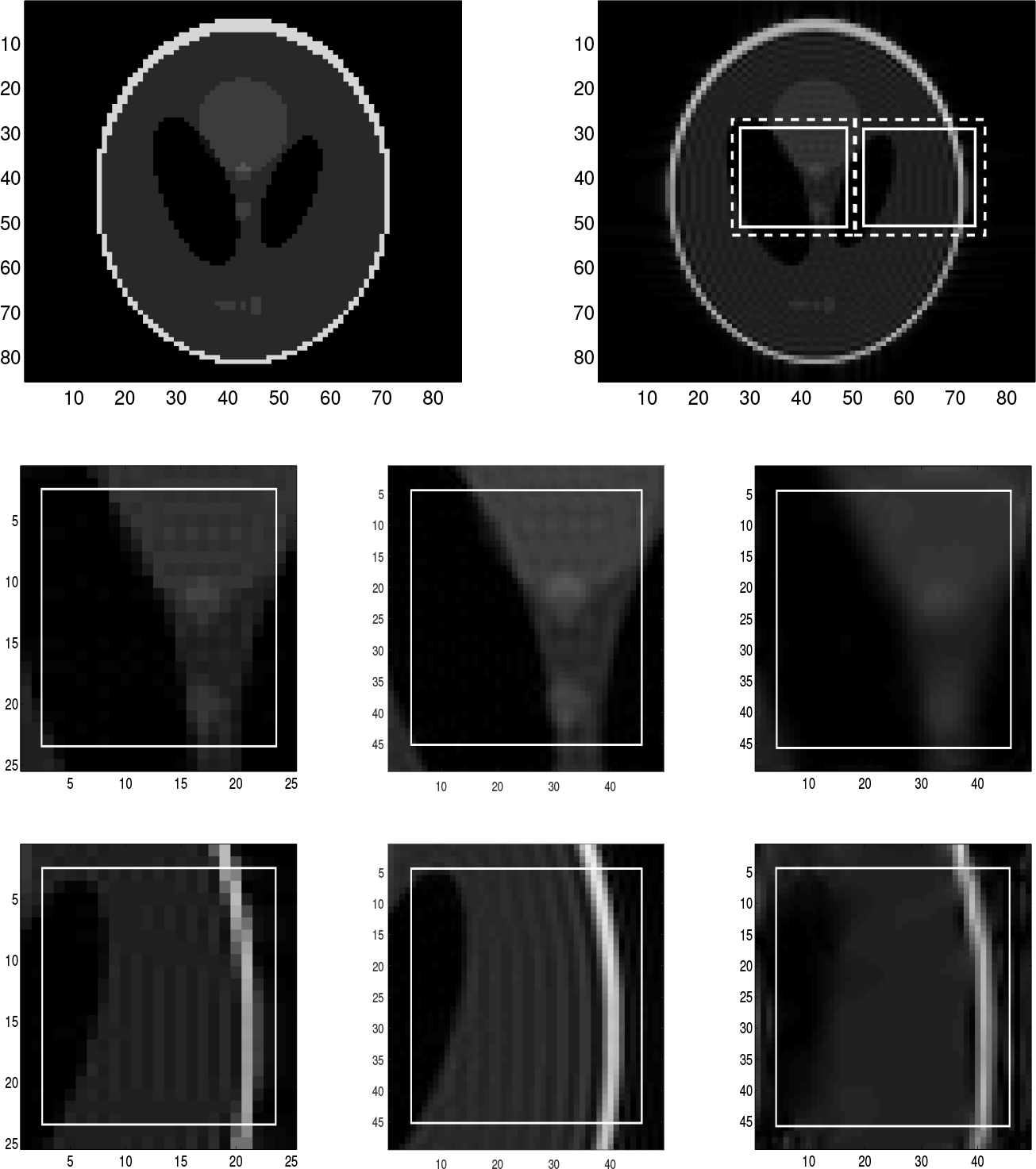
Reconstruction of Shepp-Logan phantom image, *f_N_*, by the global method. First row left: The exact Shepp-Logan phantom image on 85 × 85 grid. First row right: The Shepp-Logan phantom image, *f*_*N*_, (the Gibbs oscillations are clearly visible). The sample region 1 (left rectangle) and the sample region 2 (right rectangle). The sample regions in the rectangles with the solid line are *J* and the regions in the rectangles with dashed line are *J*_1_. Second row left: Zoomed image of the sample region 1 of *f*_*N*_. Second row middle: Double the image resolution of the subfigure in second row left by using zero-padding(for comparison purposes). Second row right right: Reconstruction of the sample region 1 with the global 2D Fourier continuation sparse PA method. Third row left: Zoomed image of the sample region 2 of *f*_*N*_. Third row middle: Double the image resolution of the subfigure in third row left by using zero-padding(for comparison purposes). Third row right: Reconstruction of the sample region 2 with the global 2D Fourier continuation sparse PA method.

We can see that for both sample regions, *J*, we eliminated Gibbs oscillations shown in the figures on the middle column by the global 2D Fourier continuation sparse PA method.

For the sample region 2, the reconstruction has distinct oscillations near the boundaries of the region *J*_1_. Since we only consider the smaller region *J*, these oscillations can be ignored. We can also see that the reconstruction on the right is blurry near the edges. As we will see in the following section, the reconstruction near edges is too smooth compared to the results by using the dimension by dimension method even though the Gibbs oscillations are much reduced.

It took about 250 seconds for the reconstruction with the global approach to be completed with Intel Core i7-3610QM and 2.30GHz.

#### Dimension by dimension method

In Fig 2 we choose the same sample regions as in Fig 1 and apply the dimension by dimension Fourier continuation sparse PA method to these two regions. We choose λ = 0.02 for both regions. The value of λ = 0.02 is also chosen by experiments. The regions in those rectangles with the solid border, *J*, are the ones where we want to find reconstruction and the whole regions, *J*_1_, are the ones with the fixed number of extra points to each boundary. The images in the left column are the zoomed images of the extended samples of the Shepp-Logan phantom. The images in the middle column are the reconstructed images of the images in the left column by applying the 1D Fourier continuation sparse PA method in *x*-direction. In these images, we eliminated the Gibbs oscillations along the *x*-direction. The images in the right column are the reconstructed images of the images in the middle column by applying the 1D Fourier continuation sparse PA method in *y*-direction. These images are the reconstructions of the images in the left column by applying the dimension by dimension Fourier continuation sparse PA method. As shown in these images, we find that the Gibbs oscillations in both *x* and *y*-directions are much reduced. By comparing those images in the first and third columns we observe that the reconstructions near both weak and strong edges are Gibbs-free and sharp. For these sample regions, the reconstructed images may have oscillations near the subdomain boundaries of the extended domain *J*_1_. Since we only consider the region *J*, these oscillations can be ignored. We also observe that the reconstructed images on the right are much sharper than the ones in Fig 1 by the global method near the edges.

**Fig 2.**
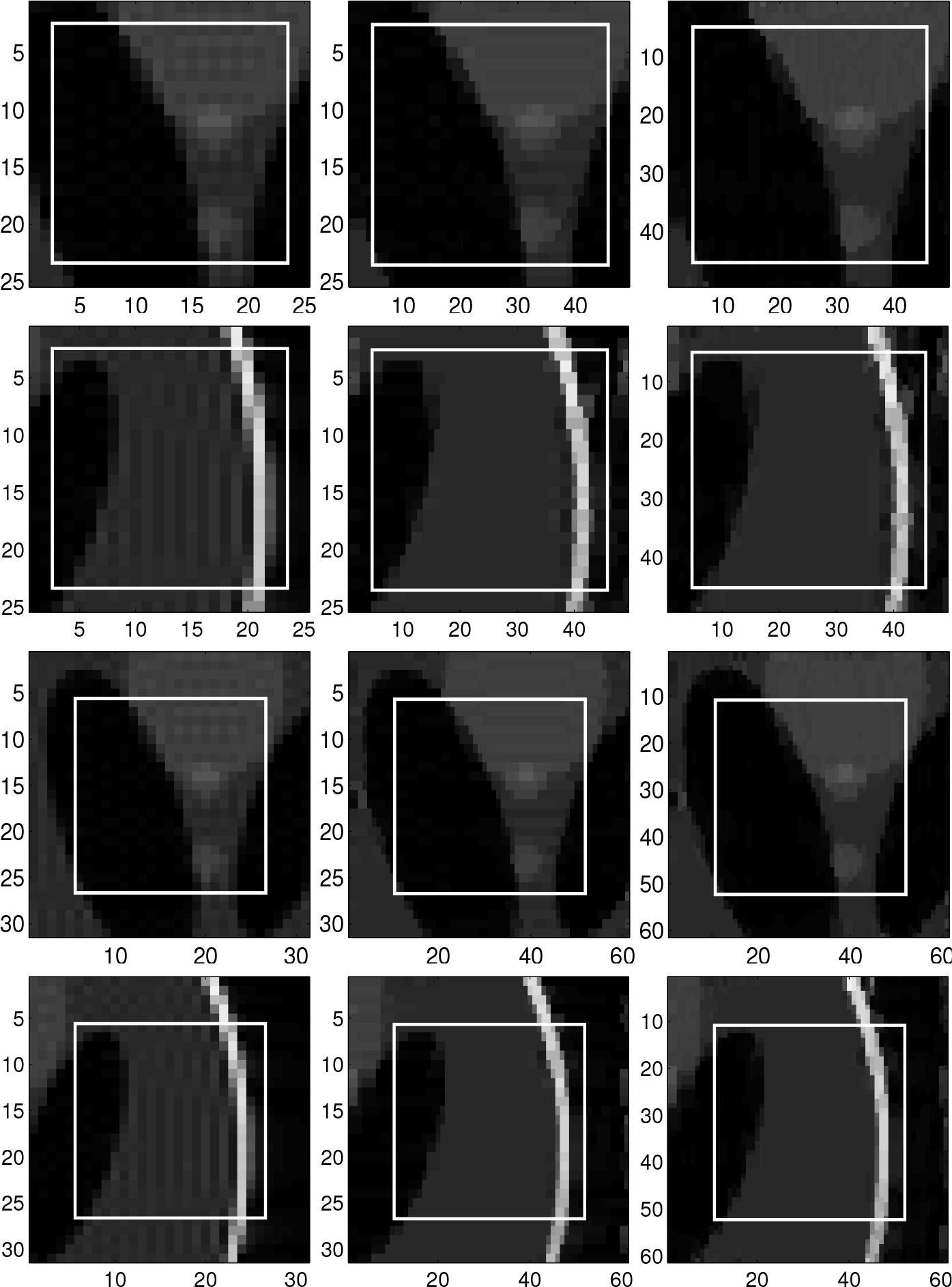
Reconstruction of Shepp-Logan phantom image, *f*_*N*_, by the dimension by dimension method. Left column: Zoomed images of the extended samples of the Shepp-Logan phantom, *f*_*N*_, on top right of Fig 1. From top to bottom, are: (1) zoomed image of the sample region 1 with two extra points added to each boundary, (2) zoomed image of the sample region 2 with two extra points added to each boundary, (3) zoomed image of the sample region 1 with five extra points added to each boundary and (4) zoomed image of the sample region 2 with five extra points added to each boundary. Middle column: Reconstruction of the image in the left column with the 1D Fourier continuation sparse PA method in *x*-direction. Right column: Reconstruction of the image in the middle column with the 1D Fourier continuation sparse PA method in *y*-direction.

Furthermore, it took about 30 seconds to complete the reconstruction in the right image in the first two rows (which have two extra points to each boundary) and about 35 seconds to complete the reconstruction of the right image in the last two rows (which have five extra points to each boundary).

In Fig 3 we split the Shepp-Logan phantom image in the top right into 5 × 5 subdomains, use the dimension by dimension Fourier continuation sparse PA method on each subdomain and finally stitch all the reconstructions from each subdomain together. By comparing the top right and bottom left images, we can see that the proposed method eliminates the Gibbs oscillations and that the error between the reconstructed image and the exact Shepp-Logan phantom image is small.

**Fig 3.**
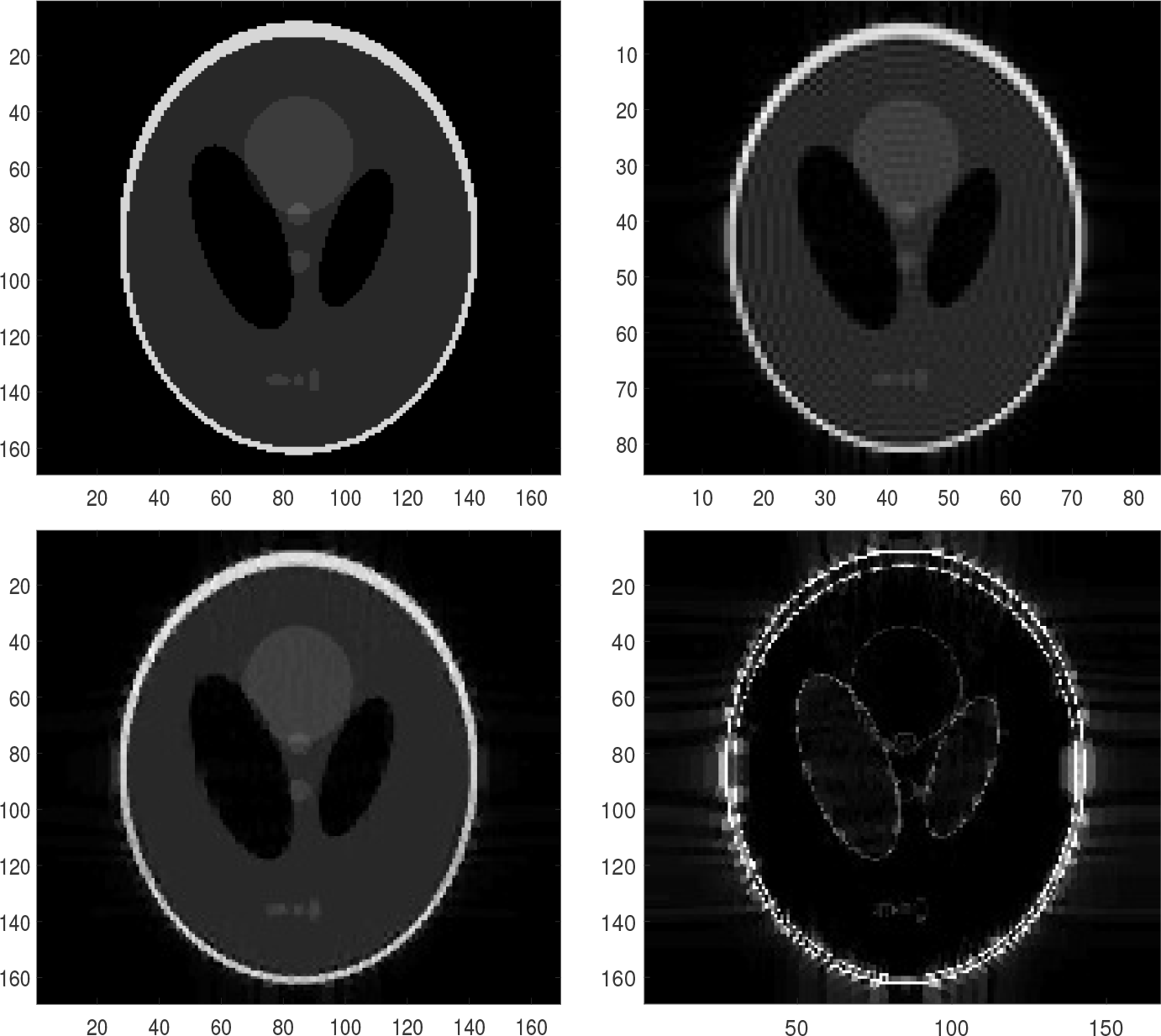
Net absolute error. Top left: The exact Shepp-Logan phantom image on 169 × 169 grid. Top right: The Shepp-Logan phantom image, the Fourier partial sum, *f_N_* with *N* = 42 (Gibbs oscillations are clearly seen). Bottom left: Stitched reconstruction constituted of 25 reconstructions from the 5 × 5 subdomains by the dimension by dimension Fourier continuation sparse PA method. For each subdomain, we have five extra points added to each boundary. Bottom right: Net absolute error image between the top left image (exact) and the reconstruction on the bottom left.

Fig 4 shows the spectrum of two Fourier coefficients of the Shepp-Logan phantom, the exact Fourier coefficients and the Fourier coefficients of the reconstruction phantom. The exact Fourier coefficients (red) are found by applying the 2D Fast Fourier transform on the 801 × 801 Shepp-Logan phantom. The Fourier coefficients of the reconstruction (blue) are found by applying the 2D Fast Fourier transform on the stitched reconstruction on 169 × 169 grid obtained by the dimension by dimension method. The left figure shows the Fourier spectrum with fixed *k_y_* = 0 for *k_x_* = −84, ···, 84. The right figure shows the Fourier spectrum with fixed *k_x_* = 0 for *k_y_* = −84, ···, 84. In Fig 4 we observe that the alteration of the Fourier coefficients after the reconstruction is not significant compared to the exact Fourier coefficients.

**Fig 4.**
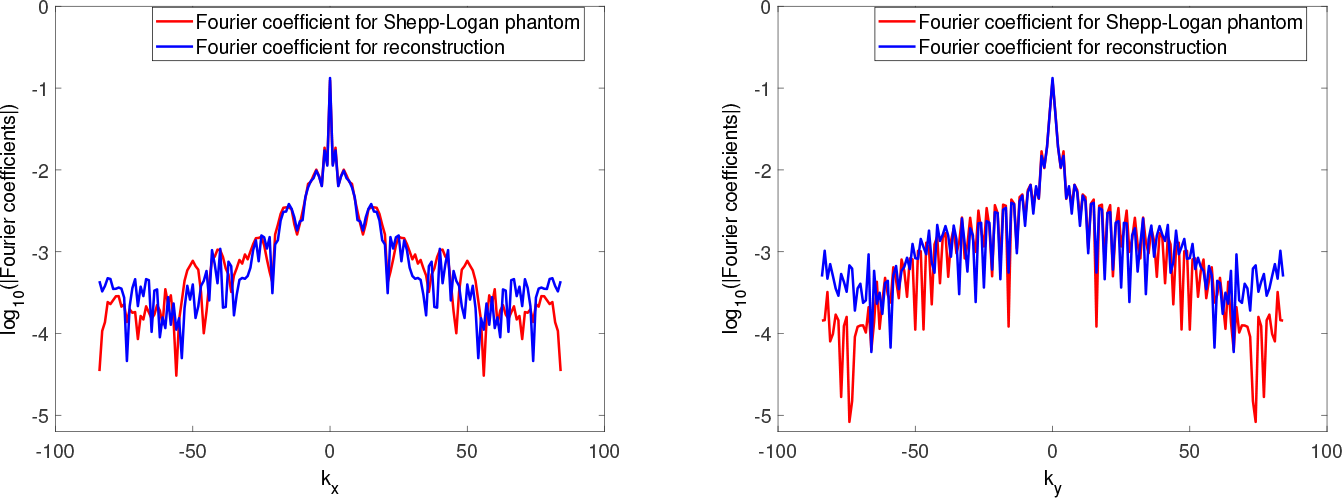
The spectrum of the exact Fourier coefficients (red) and the Fourier coefficients after stitching all local reconstructions using the dimension by dimension method (blue). Left: with the fixed *k_y_* = 0. Right: with the fixed *k_x_* = 0.

### Examle 2: MRI images

For this example, we provide the MRI image reconstructions with both the global and dimension by dimension approaches as in Example 1.

#### Global method

In Fig 5 we choose two sample regions of the MRI image and apply the global 2D Fourier continuation sparse PA method to these regions. Here we choose λ = 0. 00002 for both regions. The chosen value of λ is obtained by experiments. This value is much smaller than the one used for the Shepp-Logan phantom image with the global FC method. However, as we will see later, the lambda values are similar for both the Shepp-Logan phantom image and MRI image in this section when the dimension by dimension approach is used. This implies that the dimension by dimension approach is much more consistent than the global method in terms of choosing the optimal value of lambda. The regions in those rectangles with the solid border, *J*, are the ones where we want to find reconstructions and the regions in the rectangles with the dashed border, *J*_1_, are the extended regions of *J*. We observe that for both sample regions, *J*, the reconstructions on the right are blurry near the jumps and that there are oscillations in the smooth part as well. It took about 370 seconds to complete the reconstruction for the right image.

**Fig 5.**
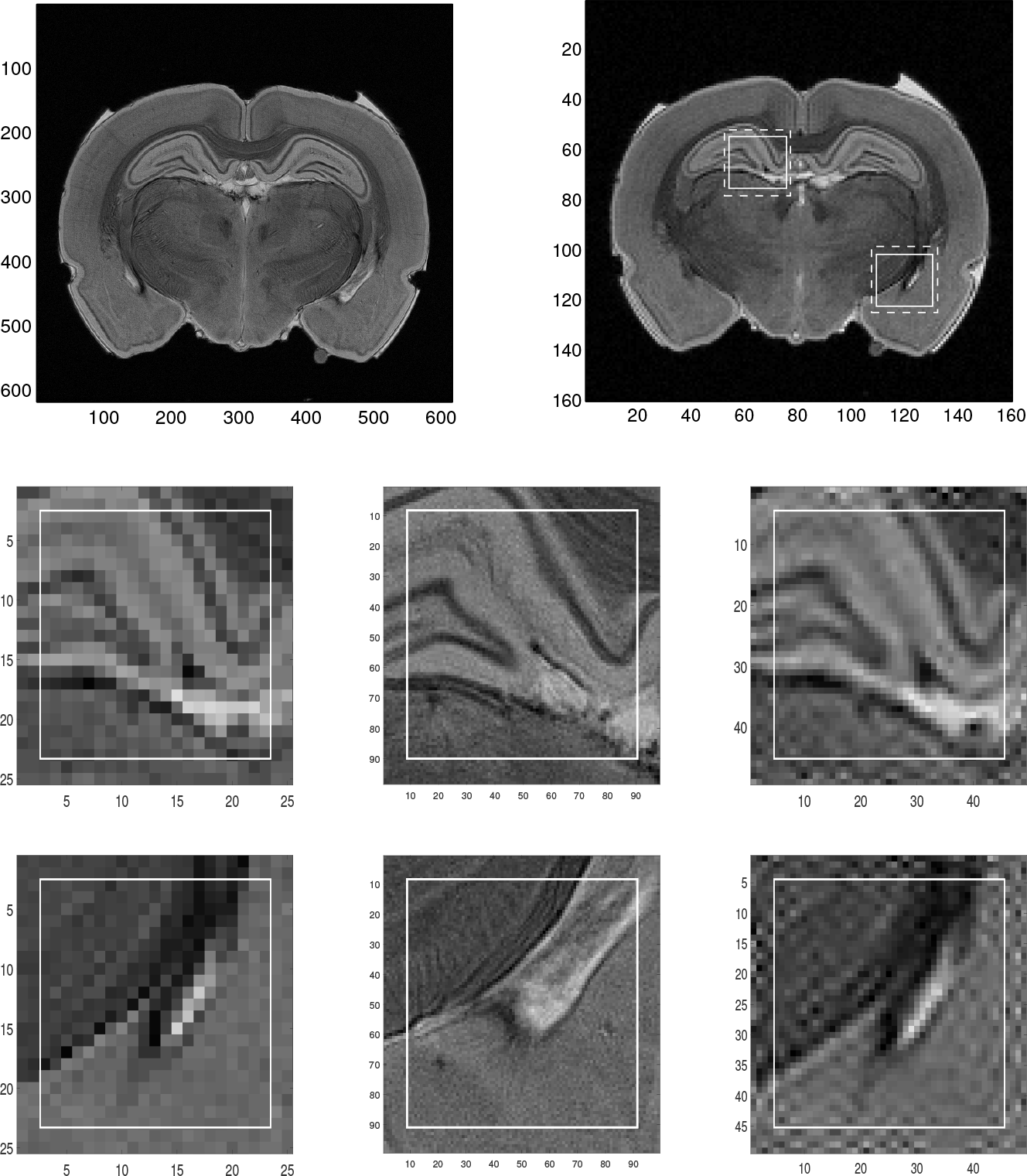
Reconstruction of the low resolution MRI image by the global method. Top left: High resolution MRI image (for comparison purposes). Top right: Low resolution MRI image. The sample regions in the rectangles with the solid line are *J* and the regions in the rectangles with the dashed line are *J*_1_. Middle left: Zoomed image of the sample region 1. Middle middle: Same zoomed sample region 1 of high resolution. Middle right: Reconstruction with the global 2D Fourier continuation sparse PA method of the sample region 1. Bottom left: Zoomed image of the sample region 2. Bottom middle: Same zoomed sample region 2 of high resolution. Bottom right: Reconstruction with the global 2D Fourier continuation sparse PA method of the sample region 2.

#### Dimension by dimension method

For the dimension by dimension method, we first show, in Table 1, the range of the value of λ that provides the best reconstruction results for the given value of *n_m_*, the number of extra points added to each boundary. *The best* reconstruction is the reconstruction that yields a sharp reconstruction near jumps while the errors in the smooth region are very small that we cannot see the difference with the bare eye. These values of λ are used for every example used in this work. This experiment suggests the range of λ values that can be used when the Fourier continuation method is applied.

**Table 1.** The best choice of λ for *n_m_*, the number of extra points added to each boundary. The value of *n* is the number of points on *x*-direction of *J*).

| $n_m$ | $\lambda$ |
| --- | --- |
| $n/10$ | $0.0175 - 0.02$ |
| $n/4$ | $0.015 - 0.0175$ |
| $n/2$ | $0.0125 - 0.015$ |

In Fig 6, 7 and 8 we chose three different sample regions, respectively, and applied the dimension by dimension Fourier continuation sparse PA method to these regions. The regions in those rectangles with the solid border, *J*, are the ones where we want to find reconstruction. The images in the left column are the zoomed images of the extended sample regions of the MRI image, which, from top to bottom, are: (1) zoomed images of the extended sample region with 2 (2 is obtained by n/10) extra points added to each boundary, (2) zoomed images of the extended sample region with 5 (5 is obtained by n/4) extra points added to each boundary, (3) zoomed images of the extended sample region with 10 (10 is obtained by n/2) extra points added to each boundary. The images in the middle column are the reconstructed images of the images in the left column with the dimension by dimension Fourier continuation sparse PA method on the region, *J*. Here we choose the value of λ according to Table 1. The images in the right column are the zoomed images of the images in the solid rectangle in the middle column. From these images we observe that if we have 5 or 10 extra points the reconstruction is sharper near the edges and less noisy in the smooth regions than we have 2 extra points. And the reconstructions with 5 and 10 extra points are not much different. For these extended sample regions, the reconstructed images may have oscillations near the boundaries of the extended region *J_1_*. Since we only need the images in the original region *J*, these oscillations can be ignored. We also observe that the reconstructed images on the right are much sharper than the ones in Fig 1 near the edges.

**Fig 6.**
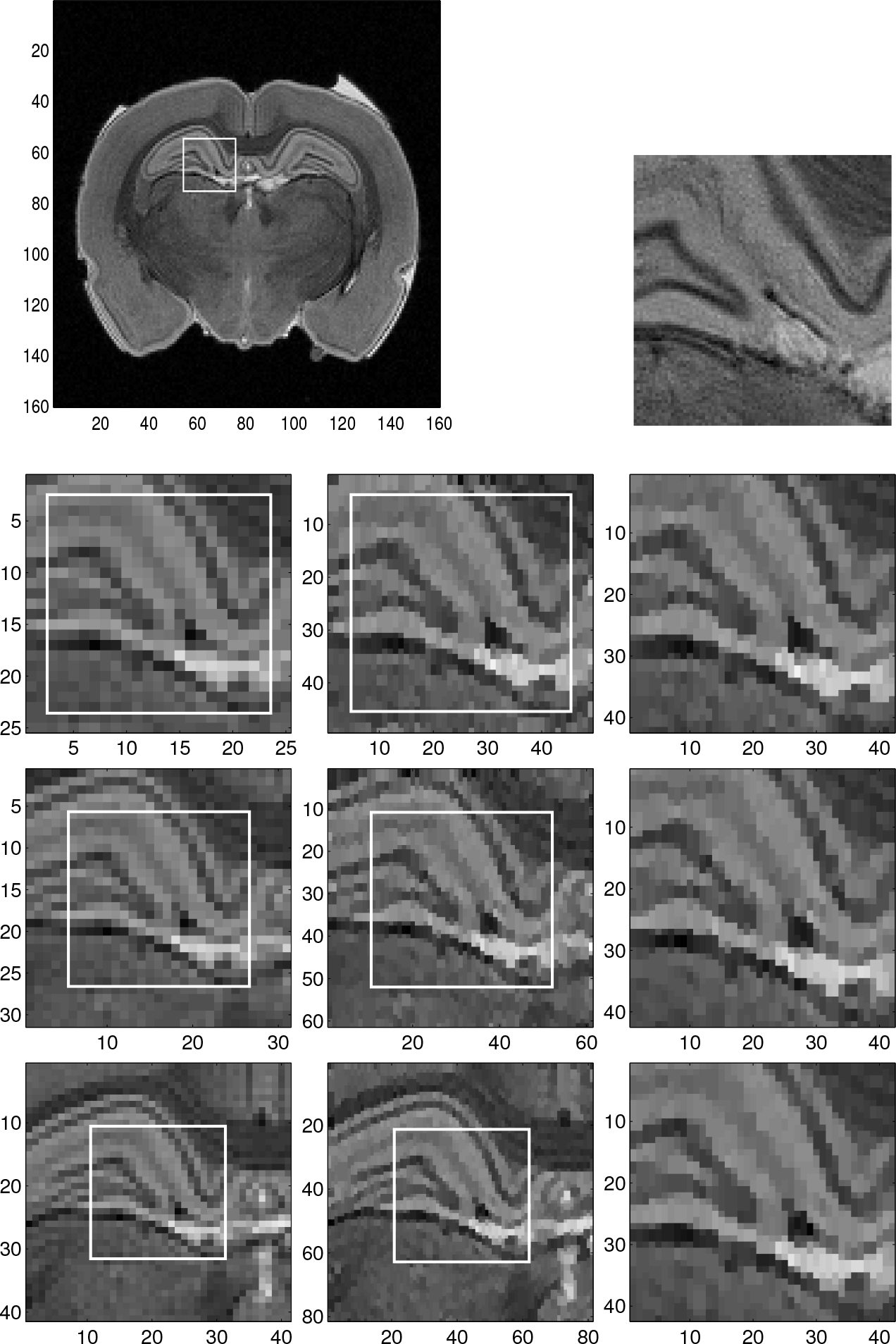
Reconstruction of the sample region 1 of the low resolution MRI image by the dimension by dimension method. Top left: Low resolution MRI image with sample region 1. Top right: Same zoomed sample region 1 of high resolution MRI image (for comparison purposes). Bottom: Left column: Zoomed images of the sample region 1 of the low resolution MRI image. Middle column: Reconstructed images with the dimension by dimension Fourier continuation sparse PA method of the sample region 1. Right column: Zoomed images of the sample region in the rectangles with the solid line in the middle column.

**Fig 7.**
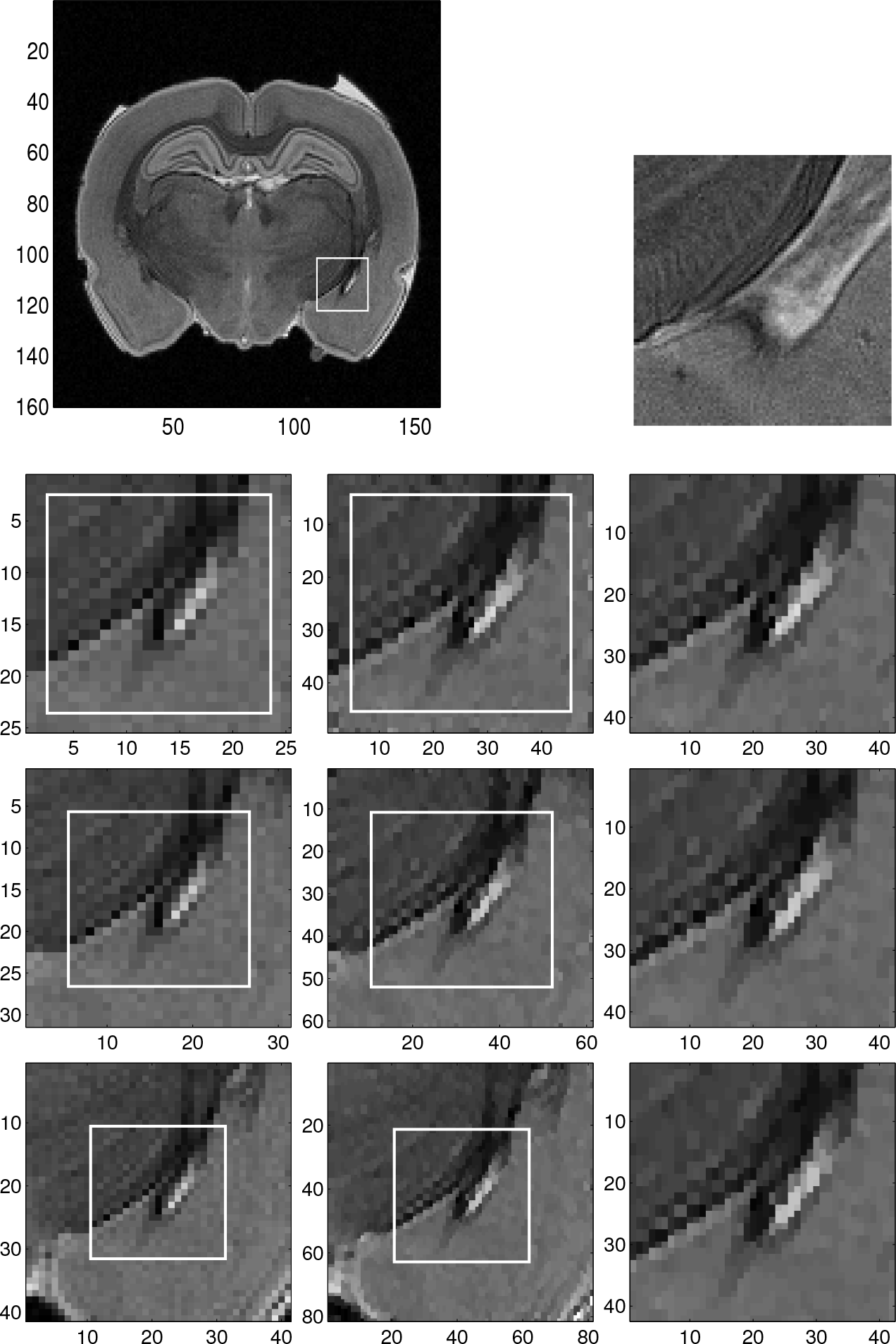
Reconstruction of the sample region 2 of the low resolution MRI image by the dimension by dimension method. Top left: Low resolution MRI image with sample region 2. Top right: Same zoomed sample region 2 of high resolution MRI image (for comparison purposes). Left column: Zoomed images of the sample region 2 of the low resolution MRI image. Middle column: Reconstructed images with the dimension by dimension Fourier continuation sparse PA method of the sample region 2. Right column: Zoomed images of the sample region in the rectangles with the solid line in the middle column.

**Fig 8.**
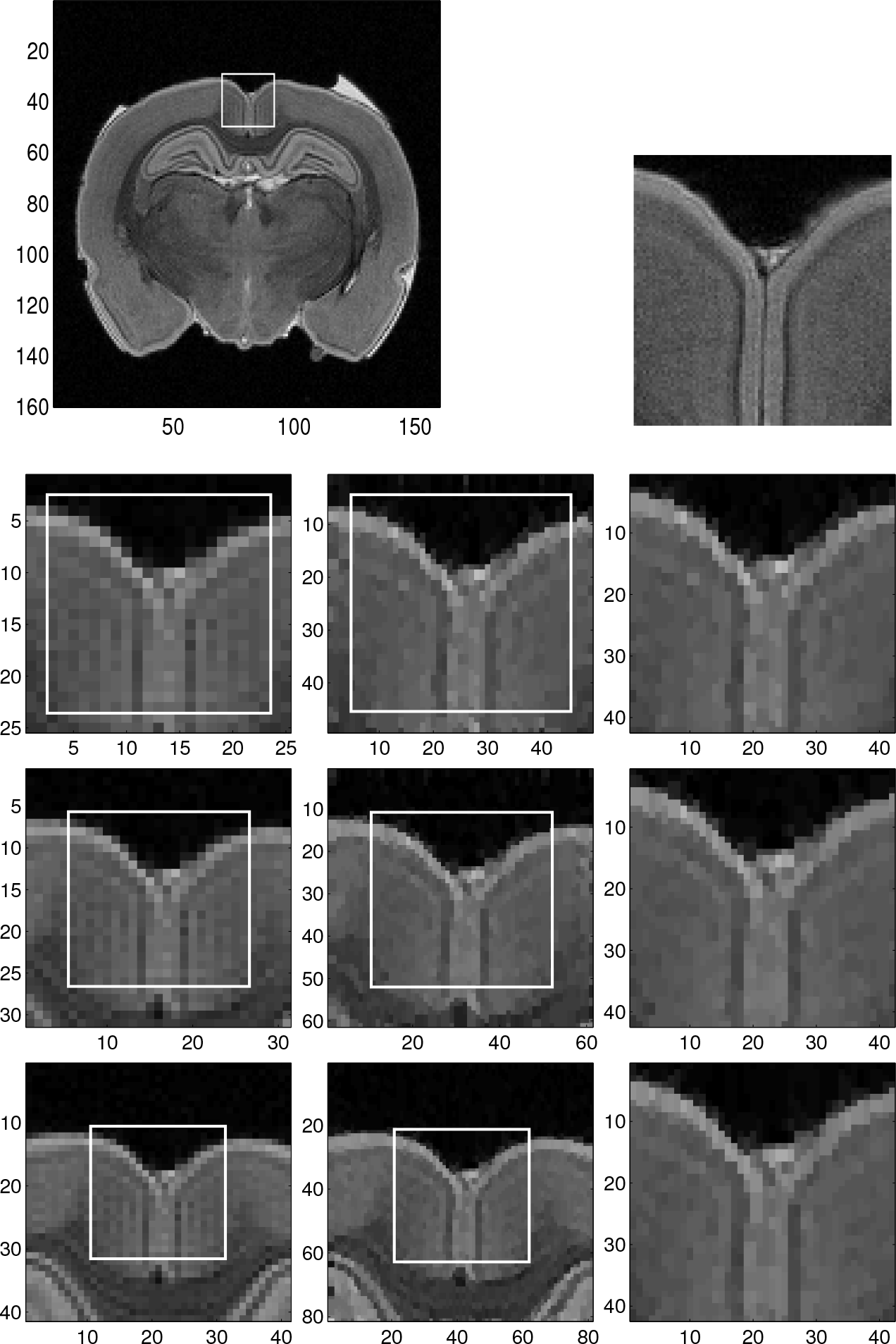
Reconstruction of the sample region 3 of the low resolution MRI image by the dimension by dimension method. Top left: Low resolution MRI image with sample region 1. Top right: Same zoomed sample region 1 of high resolution MRI image (for comparison purposes). Bottom: Left column: Zoomed images of the sample region 3 of the low resolution MRI image. Middle column: Reconstructed images with the dimension by dimension Fourier continuation sparse PA method of the sample region 3. Right column: Zoomed images of the sample region in the rectangles with the solid line in the middle column.

By comparing the reconstructions in Fig 5, 6 and 7, we find that the reconstructions with the dimension by dimension Fourier continuation sparse PA method is much better than the reconstructions obtained by the global 2D Fourier continuation sparse PA method. Furthermore the dimension by dimension Fourier continuation sparse PA method needs about 10 times less computing time than the global 2D Fourier continuation sparse PA method. Thus it is suggested that the dimension by dimension approach be used for both accuracy and computational efficiency.

In Fig 9 we split the given MRI image to 8 × 8 subdomains, use the dimension by dimension Fourier continuation sparse PA method on each subdomain and finally stitch the reconstructed subdomains together. In Fig 9 we show three reconstruction images: (1) 2 extra points (2 is obtained by n/10,n is the number of points on x direction of subdomain) added to each boundary with λ = 0.0175, (2) 5 extra points (5 is obtained by n/4) added to each boundary with λ = 0.015, (3) 10 extra points (10 is obtained by n/2) added to each boundary with λ = 0.0125. By comparing the sample regions 1 and 2 in Fig 9, we can see that in all the three reconstructions the Gibbs oscillations are eliminated. From these images we observe that if we have 5 or 10 extra points the reconstruction is sharper near the edges and less noisy in the smooth regions than 2 extra points. And the reconstructions with 5 and 10 extra points are not different. Thus it is suggested to use 5 extra points (5 is obtained by n/4) with λ = 0.015 for computational efficiency.

**Fig 9.**
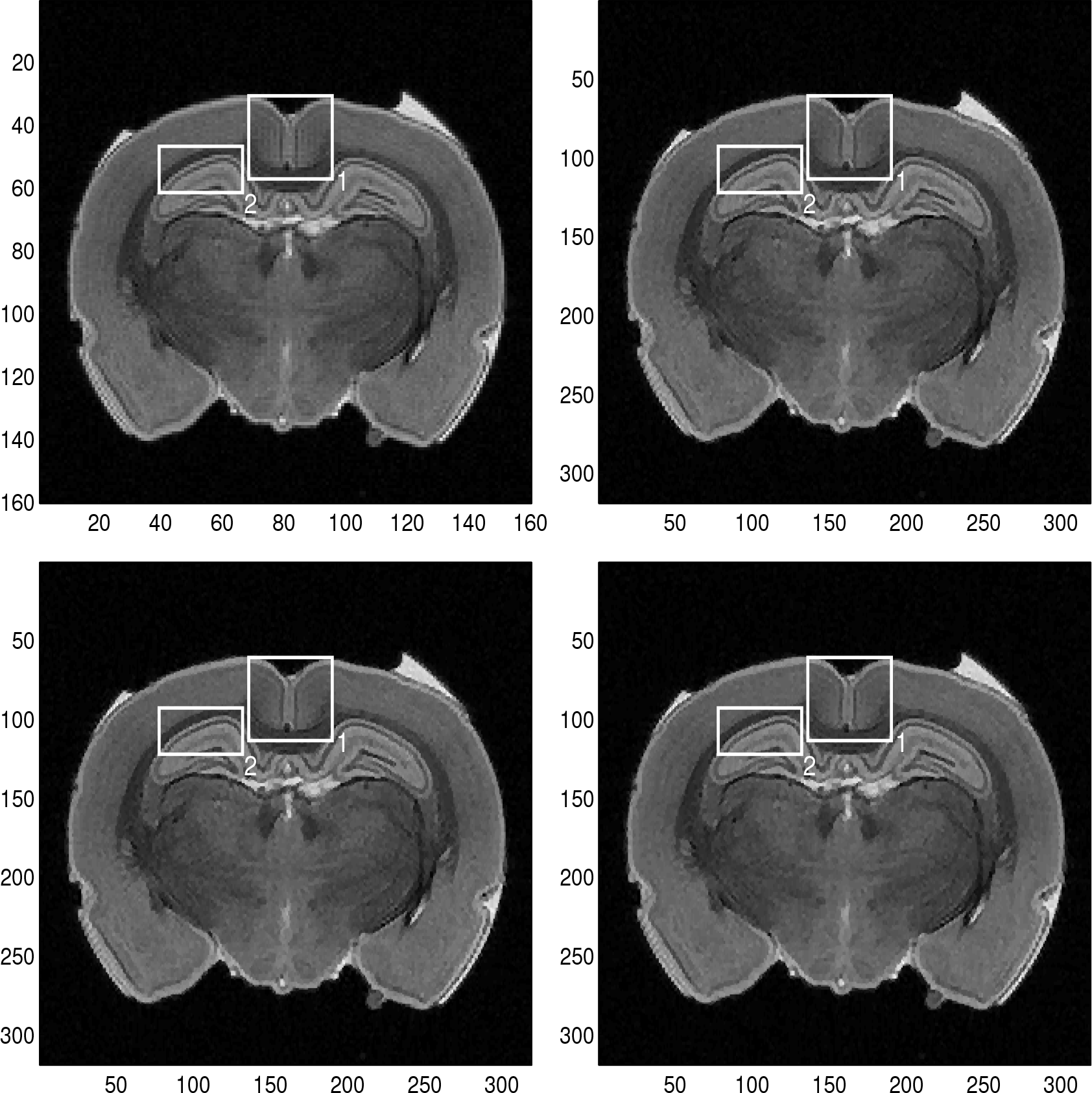
Stitched reconstruction over the whole domain. Top left: Given low resolution MRI image (for comparison purposes). Top right: Stitching the reconstructed images with the dimension by dimension Fourier continuation sparse PA method of 8 × 8 subdomains. For each subdomain, we have two extra points added to each boundary, and λ = 0.0175. Bottom left: Stitching the reconstructed images with the dimension by dimension Fourier continuation sparse PA method of 8 × 8 subdomains. For each subdomain, we have five extra points added to each boundary, and λ = 0.015. Bottom right: Stitching the reconstructed images with the dimension by dimension Fourier continuation sparse PA method of 8 × 8 subdomains. For each subdomain, we have ten extra points added to each boundary, and λ = 0.0125.
are 1 and other elements are 0.

## Conclusion

In this paper, we extend our 1D domain decomposition Fourier reconstruction method [12] to 2D image reconstruction. We propose the global 2D Fourier continuation sparse PA method and the dimension by dimension Fourier continuation sparse PA method. The global 2D Fourier continuation sparse PA method first divides the 2D image into many subdomains and applies the global 2D Fourier continuation to find the 2D periodic extension of the subdomain we are interested in. Finding new Fourier coefficients based on the periodic extension and using these we applies the 2D sparse PA method to obtain the reconstruction. The dimension by dimension Fourier continuation sparse PA method first divides the 2D image into many subdomains and in the subdomain we are interested applies the following steps on the 1D data in x- and *y*-directions separately. Applying the 1D Fourier continuation to find a periodic extension of the 1D data in x- or *y*-direction. Finding new Fourier coefficients based on the periodic extension and applying the 1D sparse PA method on these coefficients to obtain the reconstruction. By splitting the 2D image into many subdomains, we obtain sharper reconstruction near both strong and weak edges.

The numerical results in this paper show that the dimension by dimension method yields more accurate reconstruction and this method is more efficient than the global method.

The dimension by dimension Fourier continuation sparse PA method for 2D Fourier image reconstruction can be extended to three-dimensional problems by repeating the same procedure in *z*-direction after applying the method in both *x* and *y*-directions.

For our future research we can consider to improve our code and apply parallel computing to our code to make it more efficient. Second, in this paper we found our results for a range of λ, which is based on the visual inspection, we will consider how to choose the best λ numerically. And for the global method convex optimization problem Eq (16), maybe we can improve it further by the following two ways: (1) remove the penalty for high frequency contributions in *f*^*R*^ by removing first and last γ/2 columns from matrix **F** and removing the first and last γ/2 elements from **f**^R^. (2) add a binary mask B to the penalty term and get ||*B*(**Ff**^*R*^ − **f̂**)||_2_, where *B* is a (2*M* + 1) × (2*M* + γ + 1) matrix with elements in row (γ/2 + 1) to row (2*M* + γ/2 + 1) are 1 and other elements are 0.

## Acknowledgments

Research reported in this publication was funded by the National Center for Advancing Translational Sciences of the National Institutes of Health under Award Number UL1TR001412. The content is solely the responsibility of the authors and does not necessarily represent the official views of the NIH. The research by the second author was partially supported by Ajou University.

